# Response diversity is a major driver of temporal stability in complex food webs

**DOI:** 10.1101/2024.08.29.610288

**Authors:** Alain Danet, Sonia Kefi, Thomas Frederick Johnson, Andrew P Beckerman

## Abstract

Global change constitutes a major threat to biodiversity and ecosystem functioning which can materialise in the temporal stability of ecological communities. However, the majority of research on stability has focused on single trophic level communities and has not yet integrated classic theory about species richness and food web structure with more recent theory centred on response diversity and stochasticity. Using a stochastic, bioenergenetic food web model, we integrate these multiple bodies of theory to reveal that response diversity is a major driver of community stability. Moreover, our integrated theory reveals that positive stability-richness relationships emerge only in the presence of response diversity. In contrast to previous work, food web structure is only a secondary driver of community stability, but interacts with response diversity to determine the sign of the stability-richness relationship. Our study reveals identifiable pathways by which food web structure and response diversity drive community stability, and raises concerns about how the loss of response diversity (biotic homogenisation) may lead to a breakdown of community stability.

## Introduction

The question of how ecological communities respond to perturbations has been a long-standing question in ecology [1,2]. An abundant empirical and theoretical literature has established that species diversity can buffer the effect of environmental perturbations on ecological communities, thereby increasing the temporal stability of communities. This has been demonstrated in competitive assemblages [3–8] and in bi-trophic (plant-herbivores) communities [9–11].

A parallel body of theory demonstrated that community structure, and specifically food web structure, could impair or enhance temporal community stability [2,12–18], even possibly leading to a negative relationships between species richness and community stability [16,19]. Isolating the effect of species richness per se from that of food web structure is not straightforward as they tend to covary [19–21], such as the fact that species-rich communities contain more trophic levels than species-poor communities [21].

Recent empirical and theoretical research showed that species richness and food web structure both drive the temporal stability of communities [18,19]. However, the mechanisms by which species richness and food web structure drive temporal stability of complex food web communities and diversity-stability relationships has not been resolved. Advancing this knowledge requires the integration of the three different mechanisms that drive the temporal stability (hereafter “stability”) of ecological communities. In what follows, we review these mechanisms and introduce our modelling approach that integrates these three bodies of theory to ultimately provide novel predictions about when and how species richness and food web structure impair or enhance the stability of ecological communities to perturbations.

The three mechanisms are formally defined by 1) the role of species “response” diversity in buffering pertur-bations [3,15,22] which gives rise to positive stability-diversity relationships; 2) the role of trophic structure and interaction strengths [2,11–14,16,17] which can impair or enhance temporal stability; and 3) the path-ways by which they affect temporal stability, i.e average population stability, asynchrony, and compensatory dynamics portfolio effects [8,23].

Because different species have diverse niches and environmental preferences, they may react differently to the same environmental change [22,24]. This diversity of species responses to environmental change, coined as “response diversity”, has been identified as a crucial mechanism that buffers the response of ecological communities to environmental perturbations [3,7,23]. For example, in a given community, some species will respond positively and others negatively to changes in temperature, depending on their thermal niche [25]. Furthermore, it has also been shown recently that the stability of communities can be decomposed into the average population stability and the asynchrony of population fluctuations [8,23]. Average population stability quantifies the average variability in species abundances, while asynchrony refers to how much the temporal fluctuations of the different species of the community are buffering each other. Recent studies have shown that asynchrony can be further resolved into “portfolio effects”, the statistical averaging of independent species fluctuations in response to environmental perturbations, and “compensatory dynamics”, where one species’ decrease can be compensated for by the increase in another species [4,8,23].

These mechanisms have been well-established in data from single trophic level (e.g. competitive) communities, such as grasslands, butterflies, birds, and bats communities [6,26]. In these studies, response diversity increases asynchrony, and is a more important driver of community stability than population stability [26]. In these same studies, the portfolio effect was found to play a key role in the emergence of positive diversity-stability relationships in experimental grasslands [8].

Despite these data, there is limited evidence and understanding for how response diversity, population stability and asynchrony integrate to define community stability in food webs with substantial diversity and vertical complexity (i.e. trophic levels) [10,18,19,27]. Classic theory and concepts about stability-complexity [2,12], stability-structure [13,14,16,18] and stability-richness [15–17] offer some expectation. They suggest that population stability is expected to decrease with increasing interaction strength [2,13,14] and food web connectance [16] but to increase with average trophic level [19,28]. As a result, the distribution and strength of species interactions, and the average trophic level, can have strong effects on community stability as well as on stability-richness relationships [16,17]. These relationships indicate that the structure of food webs strongly influences population stability and stability-richness relationships.

The more recent theory and concepts centred on response diversity and asynchrony add further predictions [8,23]. With vertical structure and trophic interactions, consumers can offset the effects of response diversity on asynchrony, decreasing asynchrony by synchronising the dynamics of their prey [29,30], or on the contrary, increasing asynchrony by generating substantial compensatory dynamics via prey switching and trophic cascades [13,31,32].

Advancing our understanding of the drivers of temporal stability in complex communities therefore requires the integration of the previous mechanisms historically only investigated in competitive communities or food web structure. Here, we specifically investigate the effects of response diversity and food web structure on community stability and on stability-richness relationships in variable environments. The key question we ask is whether response diversity, species richness, and food web structure act additively or interactively to drive community stability.

We explore this key question by extending the bioenergetic food web model [33,34], to incorporate environmental stochasticity and response diversity [35–37]. This extension of the model allows us to interrogate the relative importance of response diversity, species richness per se, interaction strengths and trophic structure on stability.

We generated over 45000 in-silico communities with diverse richness, trophic levels and connectance, and experimentally simulated variable degrees of interaction strength, response diversity and environmental stochasticity. For each community, we evaluated the effects on stability of species richness, response diversity, community structure and stochasticity. In our analyses, stability was itself partitioned into population stability and asynchrony (i.e. portfolio and compensatory effects). Finally, we also evaluated the occurrence of negative and positive relationships between community stability and species richness.

## Methods

### Bioenergenetic model

We simulated the dynamics of complex food webs using the allometric bioenergetic model [33,34,38]. This model is widely used to simulate the dynamics of complex ecological communities because of the simplicity of its parametrisation, using the metabolic theory of ecology [39]. The model describes species biomass dynamics over time with two master equations, one for the primary producers (eq. (2)) and one for the consumers (eq.(1)):

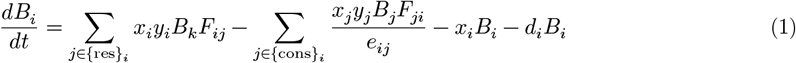

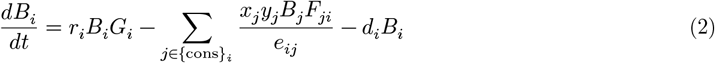

The consumers gain biomass by consuming resources (i.e. their preys) at a rate that depends on their mass specific metabolic rate (*x*_*i*_), maximum consumption rate (*y*_*i*_) and on the functional response (*F*_*ij*_) describing how the rate of consumption of a consumer *i* on a resource *j* vary with the biomass of this resource. Consumers then lose biomass by being consumed, *e*_*ij*_ being the assimilation efficiency of the consumer *j* on the resource *i*. Consumers also lose biomass over time through metabolic losses (−*x*_*i*_*B*_*i*_) and natural mortality (−*d*_*i*_*B*_*i*_). The primary producers gain biomass over time with a growth rate (*r*_*i*_, eq. (2)) and a functional response (*G*_*i*_) describing how the growth of the producers vary with their biomass. They lose biomass by being consumed by consumers and by natural mortality.

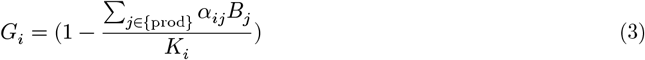

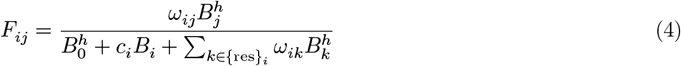

The growth of the primary producers (eq. (3)) is logistic where the growth rate is maximal and null when ∑_*j*∈{prod}_ *α*_*ij*_*B*_*j*_ are respectively close to 0 and 1. *α*_*ij*_ is the per capita effect of the producer *j* on the producer *i, K*_*i*_ being the carrying capacity for the producer *j*. The functional response of a consumer feeding on a resource (eq. (4)) depends on the relative preference of the consumer on the resource (ω_*ij*_) is limited by a half-saturation rate (*B*_0_), intraspecific interference coefficient (*c*_*i*_), and on the availability of its other resources 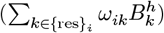. Finally, the *h* exponent controls the shape of the functional response from a saturating function (*h* = 1, Holling type II) to a sigmoid (*h* = 2, Holling type III).

### Environmental stochasticity and response diversity

We added a stochastic natural mortality rate to the species dynamics, representing a fluctuating environment. Based on the literature and previous models [35,40], we assumed that natural mortality rates scales inversely with the species body mass 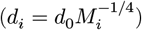, as all other physiological parameters. We used a basal natural mortality rate of *d*_0_ = 0.4, as a previous study [35].

We introduced environmental stochasticity by considering stochastic variation in mortality rates. Doing so allows to consider stochasticity without modifying species interactions themselves [35]. The stochastic part of the mortality rates (ϵ_*i,t*_) follow a normal distribution 0 centered 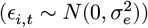 with a variance 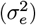. The mortality rates of each species at the time *t* equals to 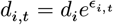, which ensured that the mortality rates never become negative [35]. We simulated ϵ using an Ornstein-Uhlenbeck process, a modified Brownian motion where the stochastic values tend to come back to the central value (i.e. ϵ = 0) such as ϵ_*t*_ varies according to this stochastic differential equation: *d*ϵ_*t*_ = (0 −ϵ_*t*_)*d*_*t*_ +σ_*e*_*dW*_*t*_, where *dW*_*t*_ is a Brownian motion. The strength of environmental stochasticity was then controlled by σ_*e*_.

Response diversity was controlled by the correlation among species stochastic mortality rates (*ρ*_*ij,i*≠*j*_) such as response diversity is maximal when the stochastic component of species mortality rates (ϵ_*i,t*_) are uncorrelated (*ρ* = 0), and response diversity is null when the stochastic component of species mortality rates are perfectly correlated [35,37,41] (*ρ* = 1). So the species mortality rates were a product of the stochastic mortality rates generated by the Ornstein-Uhlenbeck process and a variance-covariance matrix such as:

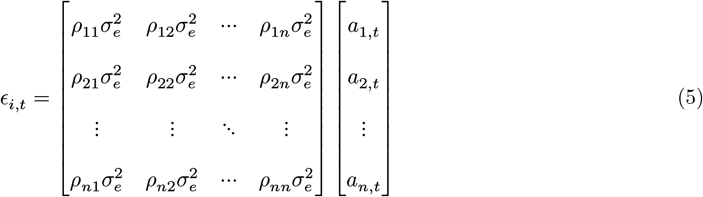

Where *a*_*i,t*_ the stochastic value generated by the Ornstein-Uhlenbeck process at time *t, ρ*_*ii*_ = 1.0 while all *ρ*_*ij,i*≠*j*_ were equal and control the level of response diversity.

### Simulation

We generated food web structure using the niche model [42] with an initial species richness from 10 to 60, and connectance from 0.02 to 0.38 (Table S1). We generated food webs without cannibalistic links and discarded food webs that contained disconnected species.

We varied the strength of environmental stochasticity by varying σ_*e*_ from 0.1 to 0.6, the range of mortality rates observed in protists [35]. We decreased response diversity by increasing *ρ* from 0 to 1 (i.e. response diversity being 1 − *ρ*) (Fig. S4).

To vary interaction strength and mortality rates (i.e. allometrically scaled mortality rate *d*) in the community, we varied the Predator-Prey Mass Ratio (PPMR) from 1 to 100 (Fig. S5b). We set *h* = 2, i.e. a functional response of type III (eq. (4)), and varied predator interference (i.e. from *c* = 0 to *c* = 1, eq. (4)), as predator interference was shown to have a tremendous role on population stability [33]. We set a global carrying capacity of *K*^′^ = 10 to ensure that consumers were not limited by the biomass input from primary producers. We standardised the carrying capacity by the number of primary producers and by the interspecific competition among producers to ensure that the effects of species richness on temporal stability are not driven by the increase in the number of primary producers, thereby trivially increasing biomass input for consumers. We then standardised *K*^′^ such as 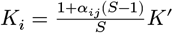 [15]. However, we found that our results were not affected when removing the standardisation of carrying capacity (Fig. S1).

We numerically solved the stochastic differential equation systems during at least 2000 timesteps and collected the last 500 timesteps, similarly to previous studies [33]. We numerically solve the system with Δ*t* = 0.1 or Δ*t* = 0.05 when instability was detected during the simulations due to stiff changes in biomass. We set extinction threshold to 10^−6^ and if an extinction happened in the last 500 timesteps, we continued to run the simulation for another 1000 timesteps until there was no extinction events during the last 500 timesteps. When we found disconnected species at the end of the simulation (i.e. a primary producer without consumer or a consumer without prey), we set their biomass to 0 and re-run the simulation for another 1000 timesteps.

Except in the case of disconnected species, we re-ran simulations with species starting biomass equal to their biomass at the last timestep of the previous simulation. Supplementary analysis showed that alternative choices of simulation processes, such as longer simulations, rebuilding food webs (i.e. resetting consumer preferences, recomputing species body mass according to the new food web) after removing disconnected species did not affect our results (Fig. S1). The food web generation and simulations were done in Julia by developing an extension of the package `EcologicalNetworksDynamics.jl` [43].

In sum, we generated 46880 simulations generated a wide range of food web structure (Median (5%, 95%), species richness: 13.0 (5.0,26.0), connectance: 0.12 (0.058,0.3), average interaction strength: 0.02(0.0031,0.071), maximum trophic level: 2.5 (2.0,3.5)) (Table S3).

### Simulation analysis

We measured food web properties that have been linked with stability in the literature. We measured total biomass, species richness, connectance, average trophic level weighted by biomass, average omnivory, and average interaction strength. food web properties were measured at each of the 500 timesteps and then averaged, except connectance and trophic level that were static. Connectance was computed as *C* = *L*/*S*(*S* − 1), *L* being the number of trophic links and *S* being the number of species in the community. Species trophic level was computed recursively from the bottom of the food web, such as the trophic level of a consumer was equal to the average trophic level of its resources added to 1, the trophic level of primary producers being set to 1 [44]. Average trophic level was then equal to: 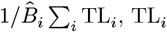 being the trophic level and 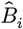 the average biomass of the species *i*. The degree of omnivory of a consumer was computed such as the sum of squares of its resource trophic levels weighted by the relative preference of the consumer for each resource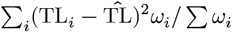 [44]. Interaction strength was computed was quantified as the biomass fluxes going from a resource to a consumer, such as: *I* = *x*_*i*_*y*_*i*_*B*_*k*_*F*_*ij*_ (eq. (1)), and averaged over time. Our way of measuring interaction strength is very similar to previous metrics used in previous studies, at the difference here that we average fluxes over time and not the maximal interaction strength [13]. Finally, the average interaction strength of a community was computed as the average interaction strength at the exclusion of null interaction strengths (i.e. excluding absence of trophic interactions).

We assessed the effects of food web structure, environmental stochasticity and response diversity on temporal stability of biomass. We measured temporal stability of biomass as the inverse of the coefficient of variation. We further partitioned stability into population stability (*S*_*pop*_) and asynchrony (*ϕ*) such as:

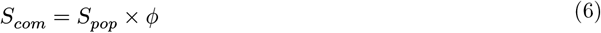

with *S*_*com*_ = *μ*_*tot*_/*σ*_*tot*_, *S*_*pop*_ = *μ*_*tot*_/ ∑_*i*_ *σ*_*i*_, *ϕ* = ∑_*i*_ *σ*_*i*_/*σ*_*tot*_; *μ*_*tot*_ and *σ*_*tot*_ being respectively average total biomass and standard deviation of total biomass.

To further get a mechanistic insight about the effects of stressors and food web structure on stability, we partitioned asynchrony into portfolio effects and compensatory dynamics stemming from species interactions [8] (*CPE*_*int*_). The measurement of portfolio effects has been a topic of debate [8,45]. Doak’s definition [4] defines portfolio effects as the product of statistical averaging effects (*SAE*) and compensatory effects arising from response diversity [4,8] (*CPE*_*env*_). It then quantities the portfolio effect (*PFE* = *SAE* × *CPE*_*env*_) emerging from independent species fluctuations, i.e. the statistical averaging effect (*SAE*), weighted by the effects of response diversity of species to environmental fluctuations, such as portfolio effects are dampened if species have correlated responses to environmental fluctuations.

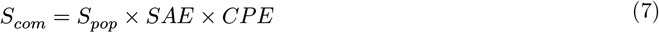

Statistical averaging effect is the share of asynchrony assuming that species are independent, meaning that the variance total of the community is equal to the sum of species variances only (i.e. species covariances are null). We can define community stability (*S*_*com,IP*_) in that scenario and then derive *SAE*.

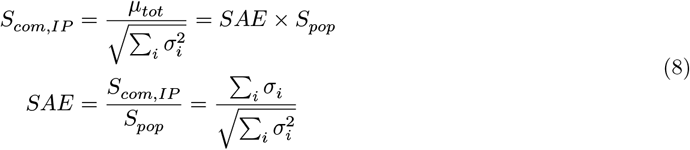

Compensatory dynamics in the food webs is then the remaining part of asynchrony:

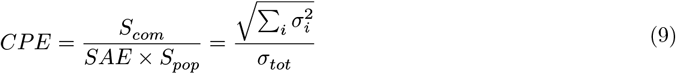

Compensatory dynamics can arise from predator-prey interactions (*CPE*_*int*_) or from differential response of species to environmental stochasticity (*CPE*_*env*_). From the deterministic and stochastic parts of mortality rates, we computed the compensatory dynamics linked to the environment such as:

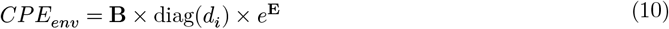

With **B** and **E** being the matrices time x species for species biomass and stochastic mortality rates respectively. Then compensatory dynamics raising from species interactions were computed as: *CPE*_*int*_ = *CPE*/*CPE*_*env*_. Finally, portfolio effects was computed as: *P F E* = *SAE*×*CPE*_*env*_ and compensatory dynamics raising from species interactions as *CPE*_*int*_. The final stability decomposition reads as follows:

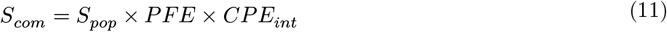

### Statistical analysis

Using a structural equation model, we disentangled the complex effects of food web structure, environmental stochasticity and response diversity on the different facets of temporal stability of biomass (i.e. population stability, asynchrony, portfolio effects and compensatory dynamic effects). We tested the effects of food web structure (average interaction strength, average trophic level, connectance) on population stability and asynchrony partitions (portfolio and compensatory effects). Grounded in theoretical ecology, we expected that higher average trophic level, lower interaction strength, lower connectance result in higher population stability and lower asynchrony [13,14,31,32]. We further added simulation parameters as control variables because higher Predator-Prey Mass Ratio and predator interference were also expected to increase population stability [33]. Average omnivory was not included because the collinearity (i.e. the VIF) with average trophic level was too high. Finally, we tested the effects of environmental stochasticity and response diversity on population stability and asynchrony partitions, although there were predicted to mainly affect population stability (i.e. by increasing amplitude of population fluctuations) and asynchrony (i.e. by generating independent population fluctuations) respectively. We completed the structural equation model with the mathematical relationship linking compensatory dynamics due to species interactions (*CPE*_*int*_) and statistical averaging effects (*SAE* × *CPE*_*env*_) to asynchrony, and from asynchrony and population stability to community stability.

Prior testing the structural equation model, we logged all the stability components to transform their relation from multiplicative to additive (eq. (7)). We ensured that all the linear models composing the structural equation model presented low multicollinearity (Variance Inflation Factor < 3, Table S4), although interaction strength and species richness were highly correlated (Fig. S5). The structural equation model using the R package `PiecewiseSEM` [46]. We computed the sum of the direct and indirect effects using `semEff` R package [47]. In the main text, we reported standardised coefficients which were obtained by scaling the coefficients by the standard deviation of the response and predictor variable. Finally, we reported only the direct standardised coefficients with an absolute value equal or above 0.05, because almost all effects were statistically significant given the high numbers of simulations.

In a second analysis, we tested how food web structure, response diversity, and environmental stochasticity modulate stability-species richness relationships. We modelled temporal stability of community biomass according to food web structure: species richness, weighted average trophic level, connectance and average interaction strength, as well as environmental stochasticity and response diversity. We further included the two-way interactions between species richness and food web structure, between species richness and response diversity, between species richness and environmental stochasticity. Finally, we added the three-way interactions between species richness, food web structure and response diversity, between species richness, food web structure and environmental stochasticity. We added predator interference, predator-prey mass ratio as control variables. We further added the ID of the food web as a random intercept. We then used the linear equation of the model and associated slope coefficients to predict the shape of the relationship between community stability and species richness according to a gradient of response diversity and of food web structure. To do so, we summed all the coefficients involving species richness in the model, i.e. the one, two, and three-way terms. The values of the other variables were set to the values indicated in the Fig. 2. The coefficients of the models are displayed in Fig. S3. The VIF were inferior to 3, indicating low multicollinearity (Table S5), we checked the distribution of the residuals (Fig. S6) using `DHARMa` R package. The model was implemented using the R package `glmmTMB` [48].

**Figure 1.**
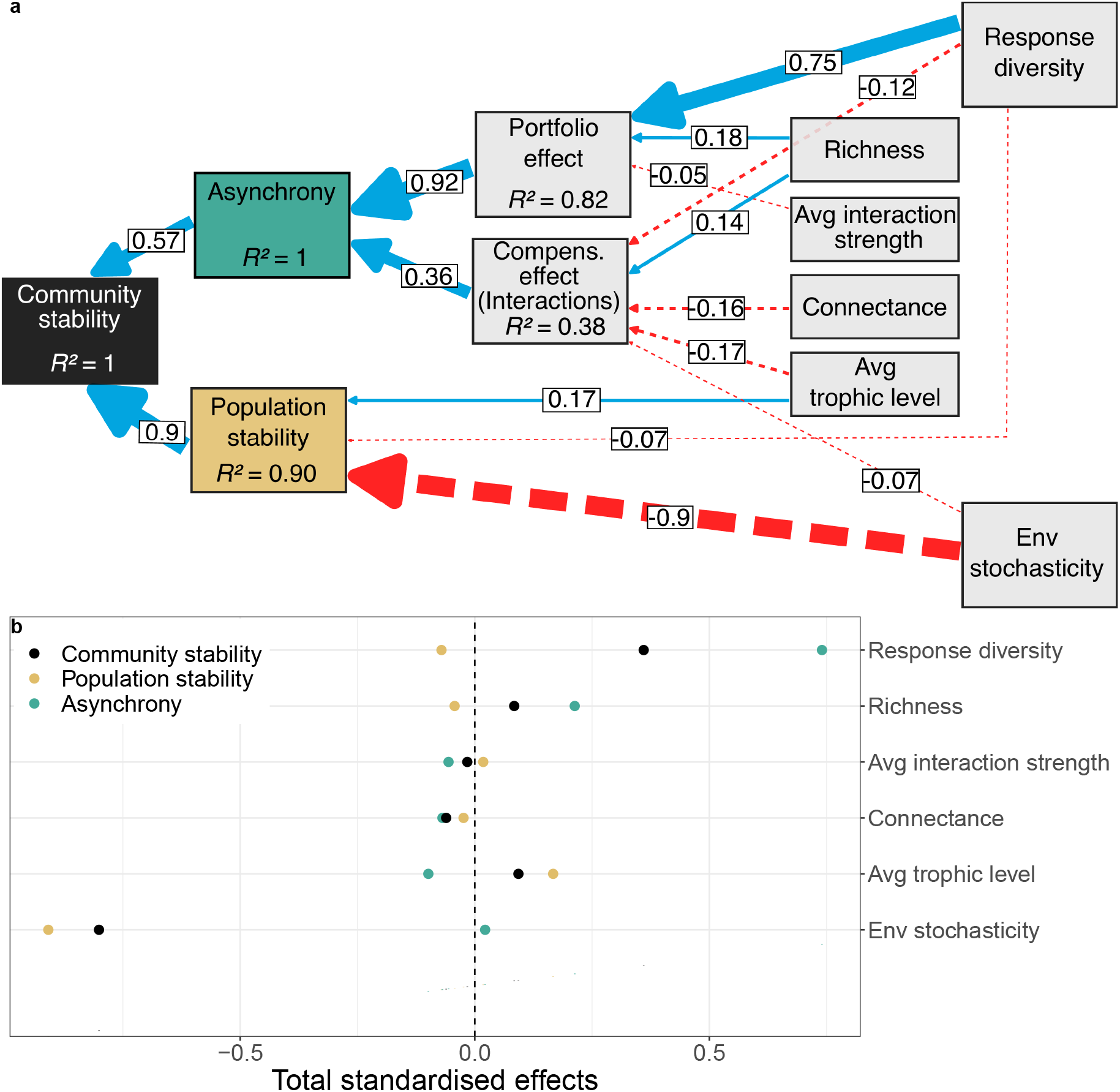
Structural Equation Model linking food web structure and stressors to the temporal stability of community biomass. a, Continuous blue arrows display positive and dashed red arrows display negative standardised effects, width of the arrows being proportional to the absolute values of the standardised effects. Only the standardised coefficients whose absolute values are superior to 0.05 are displayed. The full table of coefficients including control variables is displayed in Table S2. b, Total effects of food web structure on community stability, asynchrony, and population stability derived from the structural equation model. N = 46880 simulations.

**Figure 2.**
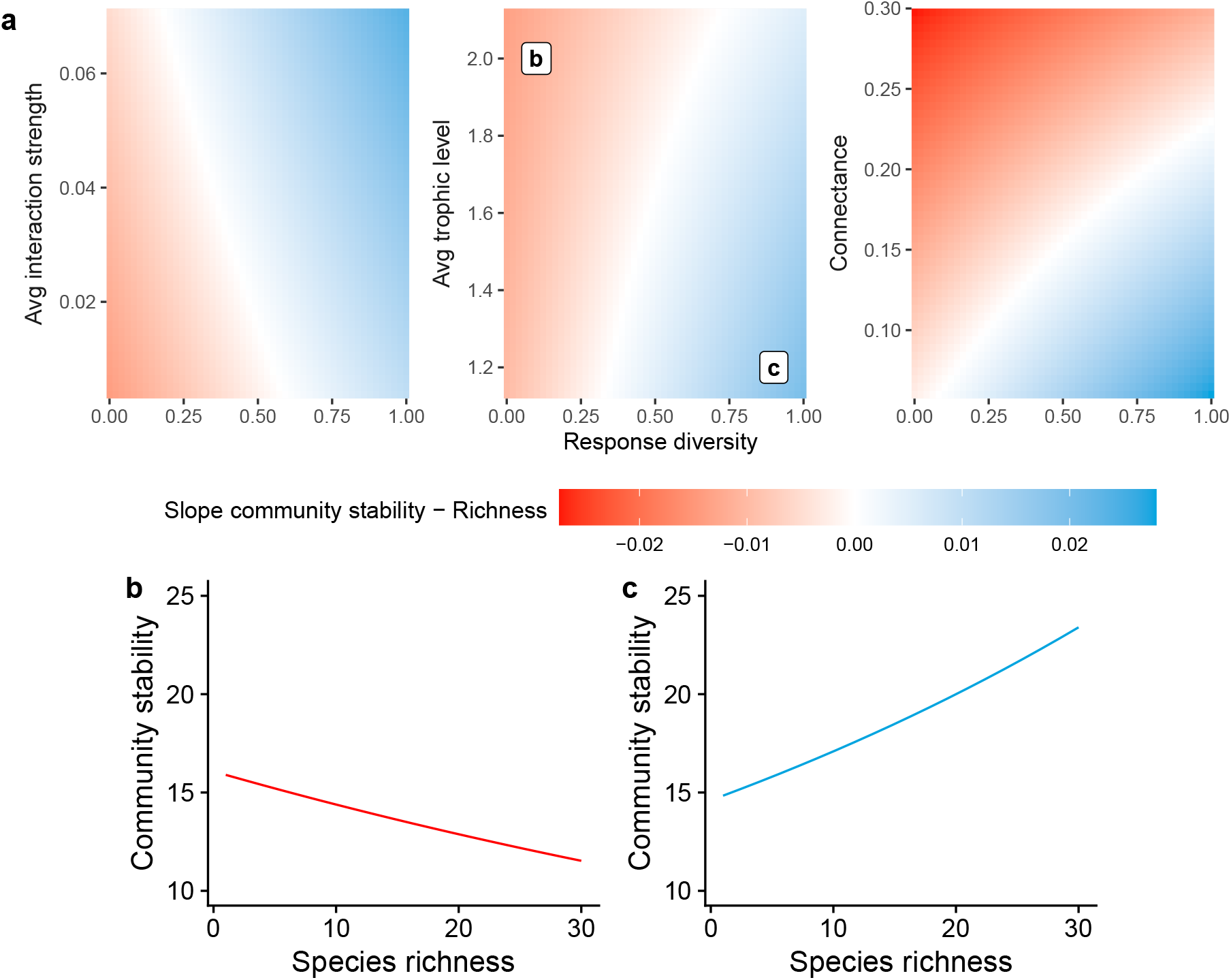
Slope of the community stability - species richness relationship based on food web structure and response diversity. a, Diagrams displaying the slope coefficients of the effect of species richness on community stability. The average values of the control variables were used to generate the predictions: connectance = 0.14, average interaction strength = 0.025, average trophic level = 1.49, predator interference = 0.5, Predator-Prey Mass Ratio = 33, and environmental stochasticity = 0.3. b, c: Examples of predictions from the model for two combinations of average trophic level and response diversity (values displayed in panel a). Lines display the mean predictions of the model. The coefficients of the linear model are displayed in Fig. S3. Relationships between population stability, asynchrony and species richness are displayed in Fig. S2.

## Results

Applying a structural equation model on the outcomes of food web simulations (see Methods) revealed that the temporal stability of community biomass is positively influenced by both population stability and asynchrony, and that population stability has twice the effect of asynchrony (resp. *r*_δ_= 0.9 and 0.57, *r*_δ_ being the standardised coefficient, Fig. 1a).

We further found that asynchrony is driven more by portfolio effects than by compensatory dynamics (resp. *r*_δ_= 0.92 and 0.36). In more details, response diversity has strong positive effects on asynchrony, mainly through portfolio effects (*r*_δ_= 0.85), at the expense of compensatory dynamics (*r*_δ_= -0.12). All other studied properties were found to have relatively much weaker effects on asynchrony. Food web structural properties – namely average trophic level, connectance and average interaction strength – have consistent negative effects on asynchrony, through both portfolio and compensatory effects (Fig. 1a). Indeed, as average trophic level and connectance increase compensatory effects are reduced (resp. *r*_δ_= -0.17 and -0.16, Fig. 1a), while average interaction strength has a small negative effect on portfolio effects (*r*_δ_= -0.05). Species richness has a positive effect on both portfolio and compensatory effects (resp. *r*_δ_= 0.18 and 0.14). Finally, as expected, environmental stochasticity has little effect on asynchrony.

Focusing on population stability, environmental stochasticity was found to have a strong negative effect on it (*r*_δ_= -0.91). All other studied properties were found to have little effect on population stability, except for the average trophic level which has a small positive effect (*r*_δ_= 0.17, Fig. 1a, but see Fig. S1) and response diversity a small negative effect (*r*_δ_= -0.07).

Deriving total effects from the sum of direct and indirect effects in the structural equation model (See Method), we showed that both environmental stochasticity and species response diversity have by far the strongest total effects on community stability (resp. *r*_δ_= -0.8 and 0.36, Fig. 1b). Species richness and average trophic level have similar small but positive total effects on community stability (*r*_δ_= 0.08), but through opposite pathways. While species richness increases community stability through a strong positive effect on asynchrony (*r*_δ_= 0.21), it also weakly decreases population stability (*r*_δ_= -0.04). In contrast, average trophic level increases community stability through strong positive effect on population stability (*r*_δ_= 0.17), but is dampened by a negative total effect on asynchrony (*r*_δ_= -0.1). Interestingly, average interaction strength also has a positive effect on population stability and a negative one on asynchrony (resp. *r*_δ_= 0.02 and -0.06). This results in a total negative effect on community stability (*r*_δ_= -0.02). Connectance, instead, has a negative effect on community stability (*r*_δ_= -0.06) through negative effects on both asynchrony and population stability (resp. *r*_δ_= -0.07 and -0.02).

We further assessed how environmental stochasticity and response diversity interacted with food web structural properties to determine the sign of the stability-richness relationship, using a linear model including main effects and all interactions among food web structural properties, response diversity and environmental stochasticity (see Methods). We found that there are strong interactions between response diversity and food web structure, which jointly determine the sign of the stability-richness relationship.

In the absence of response diversity, we found only negative stability-richness relationships, regardless of the food web structure (Fig. 2a). Response diversity enables the rise of positive stability-richness relationships, enhanced by average interaction strength and dampened by average trophic level and connectance. Higher interaction strength leads to positive stability-richness relationships for lower values of response diversity, and interaction strength enhances asynchrony-richness relationships (Fig. S2), so that the stronger positive stability-richness relationships are found at both high interaction strength and high response diversity levels. In contrast, higher average trophic level and connectance both lead to negative stability-richness relationships, because they drive more negative population stability-richness relationships and weaker positive asynchrony-richness relationships (Fig. S2). Overall, our results highlight a strong contrast between the relatively small effects of food web structural properties on overall community stability (Fig. 1b) but their strong effects on the slope of the stability-richness relationship (Fig. 2a).

## Discussion

With the projected changes in climate and land use, and the associated changes in community composition (e.g. loss of diversity, change in structure, homogenisation), it is important to understand the mechanisms which allow ecological communities to remain stable despite perturbations. Recent research has focused on the average stability of the populations composing the community and the asynchrony of the population fluctuations on the other hand [8,23]. Much of this previous work has been doneis in simplified assemblages, such as single trophic level communities, and in this contexthere community stability has been shown to be driven by asynchrony in species fluctuations, rather than by population stability [26]. However, whether this holds for more complex trophically structured ecological communities, such as food webs, remains largely unknown. Here, we tackled this question using simulations of complex, stochastic food-web dynamics via a bioenergetic model.

Our model results show that, in complex food-webs, community stability is more driven by population stability than by asynchrony, in contrast with what was observed in single trophic level communities. OurThe study further reveals that the main drivers of community stability are environmental stochasticity and response diversity, respectively acting on population stability and asynchrony. Despite previous work in simple communities suggesting that food-web structural properties drive stability, in our complex system model, these properties have much weaker effects, a result consistent across methodological choices (Fig. S1). However, structural properties were found to play an important role in the sign of the stability-richness relationship, which is strongly mediated by the interaction between food-web structure and response diversity. These results highlight the importance of response diversity to improve our understanding of the stability of structured ecological communities in variable environments.

### Response diversity and food-web structure drive community stability

We found environmental stochasticity and response diversity to be the main overall drivers of community stability. While environmental stochasticity decreases population stability, response diversity generates asynchrony, thereby buffering environmental stochasticity. This result is in accordance with previous theoretical and empirical findings in competitive communities [6,7,15,16,23,49]. In turn, response diversity was found to operate on asynchrony mainly through portfolio effects in agreement with empirical findings in competitive assemblages [7,8]. This suggests that response diversity generates asynchrony mainly by the statistical averaging of random independent species fluctuations [4].

A consequence of the large effect of response diversity is that asynchrony in species fluctuations is driven more by the statistical averaging of independent species fluctuations (portfolio effects) than by compensatory dynamics emerging from species interactions. This finding is also coherent with previous studies in competitive communities [7,8], but it was less expected to happen in food-webs. Trophic interactions are indeed expected to result in more compensatory dynamics resulting from oscillations between predators and prey [50] or prey switching from predators [13,31]. However, pioneering theoretical studies have shown that environmental stochasticity can dampen prey switching [35] and that trophic cascades between trophic levels in species rich food webs are weak compared to food chains and species poor food web modules [32]. So, the question of the importance of compensatory dynamics in complex food webs remains an open and interesting avenue for future research.

In contrast to empirical evidence in competitive communities [26], population stability (not asynchrony) was found to be the dominant driver of community stability in our food web simulations. These results are in agreement with previous findings in empirical food webs [19] and food web models [18], which suggest fundamental differences between competitive communities and food webs in the relative contributions of population stability and asynchrony to community stability. Our findings suggest that the presence of trophic links contributes to synchronise the species of the community because we found that the food web metrics studied – namely average trophic level, connectance, and average interaction strength – all decrease asynchrony. These results are in line with empirical evidence that predators can synchronise their prey [29,30], and that species pairs involved in trophic interactions are more synchronous than those that are not [19]. A complementary explanation for the relatively lower importance of asynchrony in food webs (compared to single trophic level communities) could be related to the biomass structure of food webs. Food-webs typically include a larger range of body masses than single trophic level assemblages [51]. Such highly uneven biomass distribution can translate in uneven species temporal variability, thereby creating uneven species contributions to asynchrony [23] and limiting portfolio effects and compensatory dynamics in species-rich communities [8].

Investigating in more detail the drivers of community stability, our results suggest that species richness and the network structural properties investigated only have a weak effect on overall community stability. We found that higher average trophic level leads to higher community stability by increasing average population stability, confirming previous theoretical [18,28,33] and empirical [19] results. However, this effect disappeared when we simulated community dynamics with equal mortality rates for all species instead of allometric ones (Fig. S1). This suggests that the higher stability of food webs with higher average trophic levels stems more from their lower mortality rates rather than from their trophic role directly. The overall effect of connectance on overall community stability was found to be quite low, in agreement with previous empirical and theoretical findings [18,19]. Similarly, we found very low effects of the average interaction strength on community stability, which is in apparent disagreement with the predominant role of interaction strength and connectance on stability [2]. We found that average interaction strength had a weak positive effect on population stability, in confirmation with previous theoretical findings aligning with our model assumption that there was no asymmetry of consumer preferences for their prey [52].

### Stability-richness relationships depend on response diversity and food web structure

Our results also allow us to revisit the old question of the relationship between the complexity of ecological communities and their stability [2,12]. We found that response diversity enables the rise of positive stability-richness relationships, enhanced by average interaction strength and dampened by average trophic level and connectance. In the absence of response diversity, only negative relationships between community stability and species richness are observed. This reinforces the idea that increases in species richness increase community stability only if it adds species that respond differently to environmental perturbations, i.e. if there is response diversity [6,16]. Indeed, a community with numerous species which respond differently to environmental change is more likely to buffer a wide array of perturbations [3,15]. Despite its relatively weaker effect on community stability (than response diversity and environmental stochasticity), food web structure interacts strongly with response diversity to determine the sign of stability-richness relationships. Surprisingly, we find that higher average interaction strength enhances positive stability-richness relationships by enhancing asynchrony-richness relationships. Conversely, higher average trophic level and higher connectance both led to negative stability-richness relationships by dampening asynchrony-richness relationships. Our results resonate with findings that some community structure can lead to negative stability-richness relationships, as previously reported in freshwater food webs [19]. Response diversity might therefore be one of the mechanisms that explains why positive relationships between stability and species richness are so prevalent in empirical settings [22,23], while food web theoretical models that do not often include environmental stochasticity and response diversity typically find negative complexity-stability relationships [36].

In conclusion, our study highlights the complex but consistent effects of response diversity and food web structure on the temporal stability of species rich communities. At the core of the insurance hypothesis [3,45], our results suggest that environmental stochasticity and response diversity are the main mechanisms driving community stability, while food web structural properties are only mediating them. This aligns with the idea that, in species-rich communities, community structure might be secondary in understanding the generic effects of biodiversity on ecosystem functioning [15,53]. Our results further add to mounting evidence that population stability is a stronger driver of community stability than asynchrony in food webs, in contrast with what was observed in competitive communities. The fact that environmental stochasticity was found to be the main driver of overall community stability raises concerns about the consequences of the current rise in the frequency and severity of environmental extreme events for ecological communities [54,55]; indeed our results suggest that these changes are unlikely to be compensated by biotic mechanisms alone. Documenting how rapid biodiversity changes [56,57] driven by human pressures [58] – and in particular the reported homogenisation [59,60] in community composition – affect response diversity will be key to accurately predict how it will affect, in turn, the stability of communities.

Looking forward, we emphasise promising perspectives in further understanding the temporal stability of ecological communities. Our study brings evidence that asynchrony is predominantly driven by portfolio effects as previously reported in plant communities; However, we expect that some ecological contexts could result in the dominance of compensatory dynamics over portfolio effects. Such context includes the structure of response diversity, i.e. different groups of species that respond differently to environmental stochasticity but that respond similarly within their group, are expected to give more importance to compensatory dynamics due to environmental fluctuations. The knowledge of response diversity structure in natural settings is expected to rise with the methodological progress in measuring response diversity [24,61]. Another context to explore is the effect of the distribution of interaction strength in food webs. Implementing asymmetric distributions of consumer preferences for their prey could also increase the importance of compensatory dynamics due to species interactions as strong asymmetry in interaction strength can enhance prey switching behaviour [13,31].

## Data accessibility statement

The code used to run the simulations, analyse them and to write the manuscript is archived anonymously on Zenodo: https://zenodo.org/records/14333268.

