## Supplementary material for "Response diversity is a major driver of temporal stability in complex food webs"

### Supplementary information

#### Figures

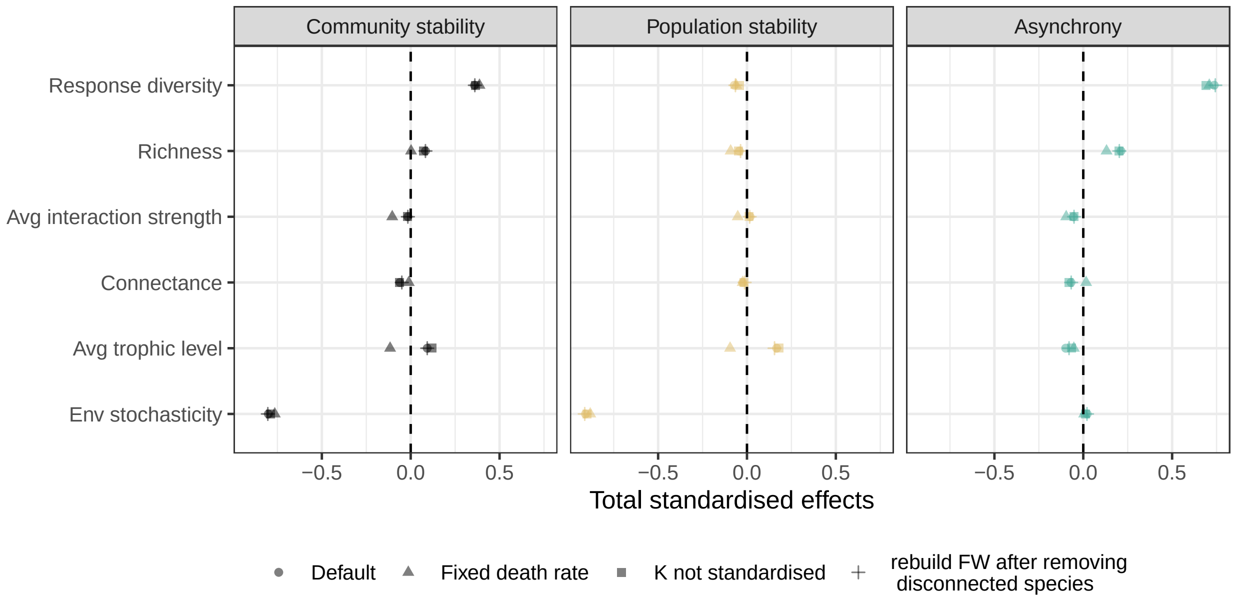


Figure S1: Total standardised effects for the sem displayed in main text according to different simulation set-ups. Default: simulations used in main text; Fixed death rate: non-allometric death rate, meaning that all species have the same death rate, here $d = 0.1$; K not standardised: carrying capacity not standardised; rebuild FW after removing disconnected species: instead of setting starting biomass of disconnected species to 0, we removed them from the food-webs, and then recompute species trophic level, species bodymass and then their metabolic rates, prey preferences, producer carrying capacity.

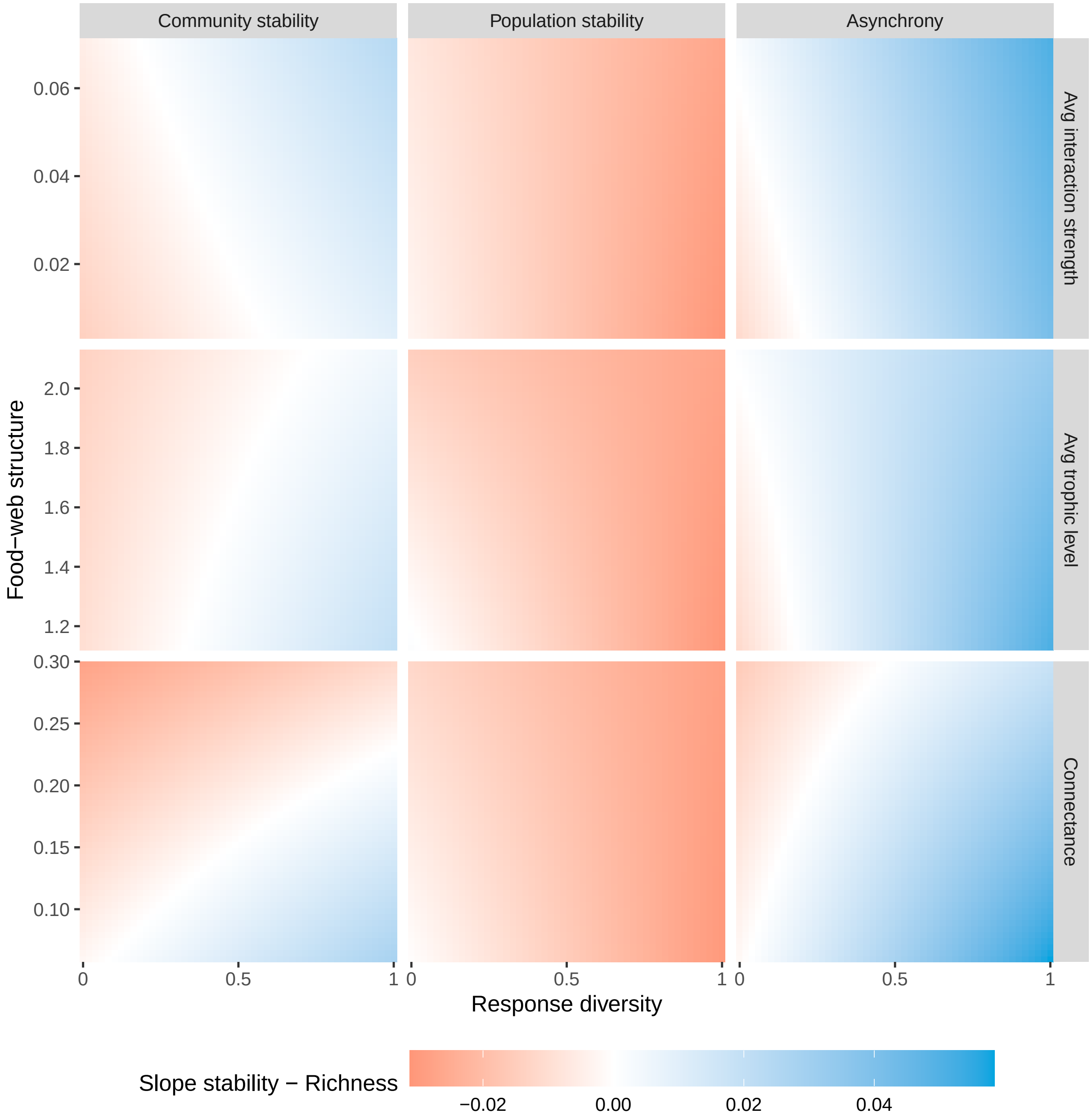


Figure S2: Prediction of the statistical model linking the temporal stability of community biomass and species richness. Compared to Fig.3, we added the prediction of the slope of the relation population stability-species richness and asynchrony-species richness. Default parameter values: connectance = 0.14, average interaction strength = 0.025, average trophic level = 1.49, predator interference = 0.5, Predator-Prey Mass Ratio = 33, and environmental stochasticity = 0.3.

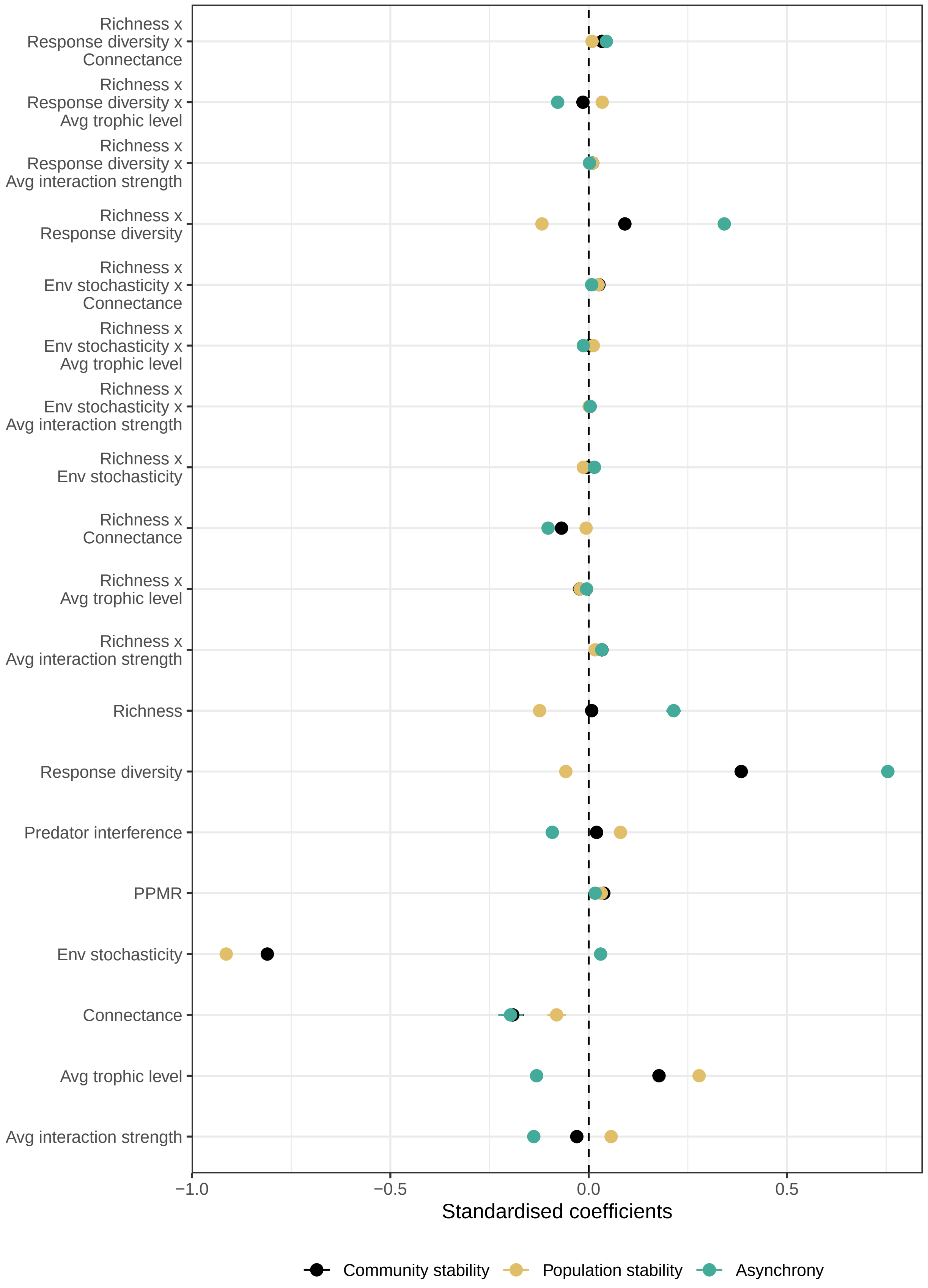




Figure S3: Standardised coefficients for the linear model predicting the relationship between community stability and species richness (Fig. 3).

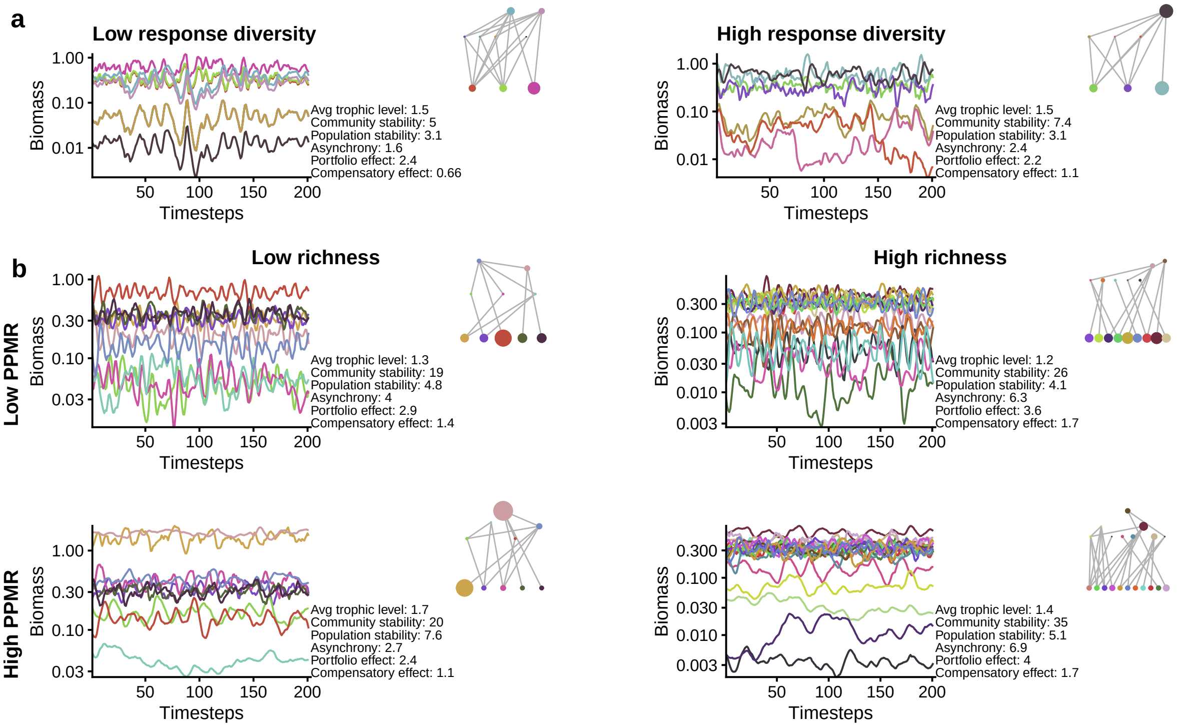


Figure S4: Examples of stochastic food-web simulations. a, Simulations with low and high response diversity. b, simulations with low and high predator-prey mass ratio (upper and bottom panels), low and high species richness (left and right panels). The food-webs are displayed on the side with the node size being proportional to the average species biomass across the simulation. We also display the food-web structure and the stability metrics. Across all simulations, $h = 2$, $\sigma_e = .3$. Low and high response diversity were respectively $\rho = 1$ and $\rho = 0$. Low and high PPMR were respectively 1 and 100.

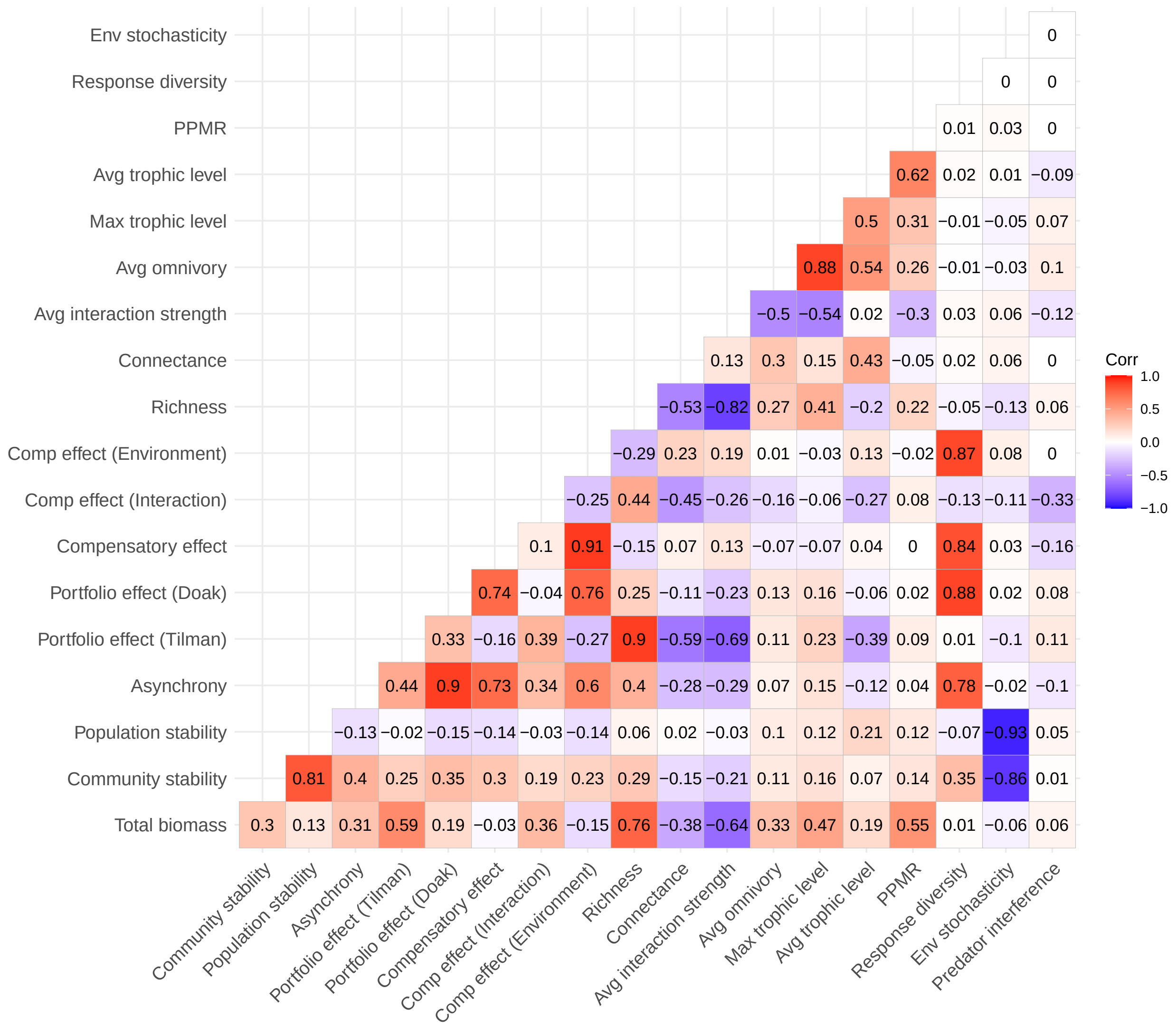


Figure S5: Spearman's correlation coeffients among stability components, food-web structure, and model parameters.

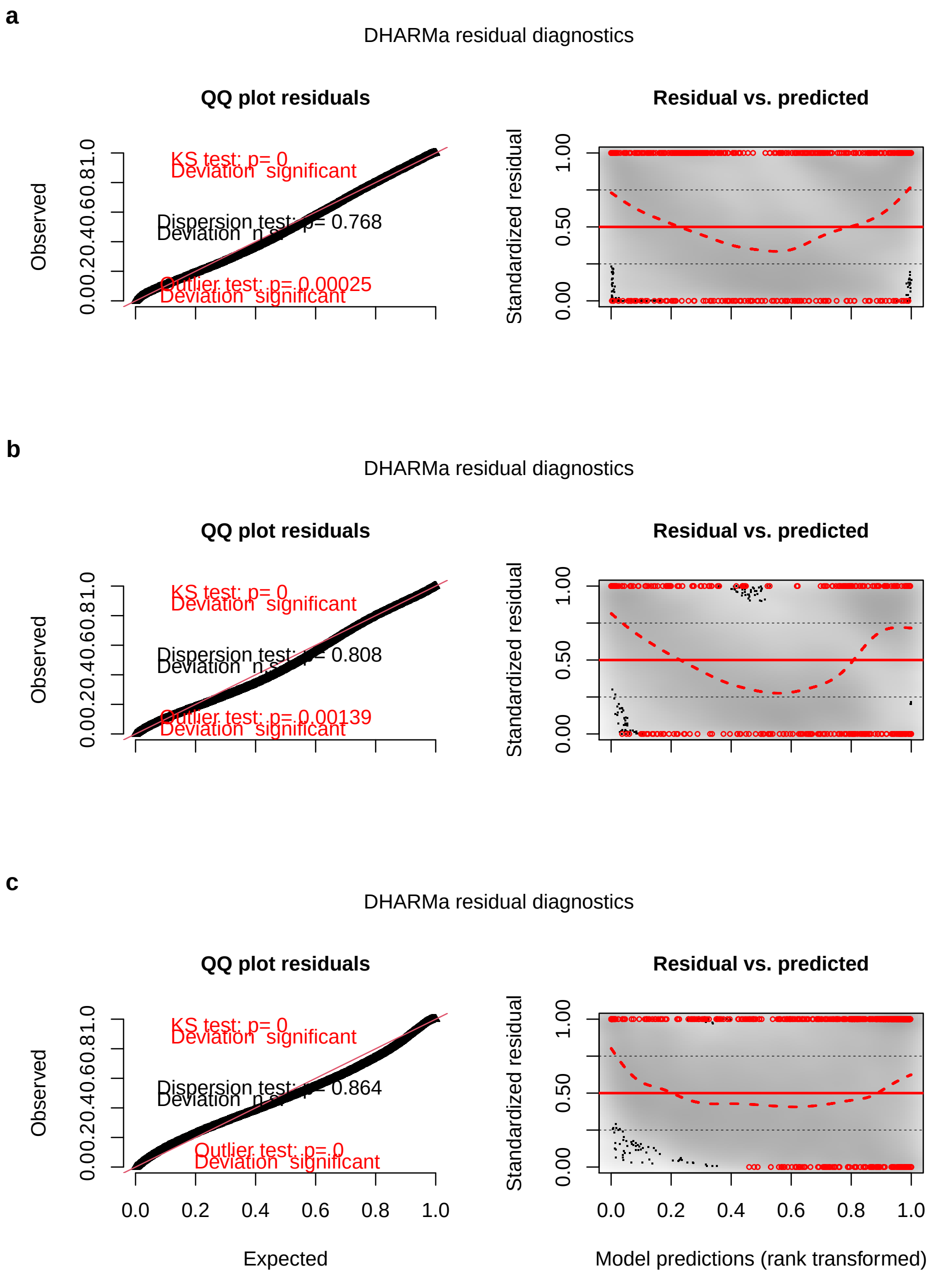


Figure S6: Distribution of residuals for the linear models predicting the relationship between community stability and species richness (Fig. 3). a,b,c for respectively community stability, population stabilty and asynchrony. Left panels display the distribution of the quantiles, and indicate that residuals are quite well distributed. Right panels display the scaled residuals versus the predicted values. The slight U-shape might indicate the lack of a polynomial term in the model, but overall the model looks to have appropriate fitting.

### Tables

Table S1: Parameter values used in the simulation experiment. N = 46880 simulations. $Z \times \rho \times \sigma_e \times c \times S = 1440$ parameter combinations. We generated 39 food-webs of varying connectance for each parameter combination.

| Parameter | Values |
| --- | --- |
| Hill exponent ($h$) | 2 |
| Half-saturation rate ($B_0$) | 0.5 |
| Producer maximum growth rate ($r$) | 1 |
| Carrying capacity ($K$) | 10 |
| Producer inter-competition ($\alpha_{ij}$) | 0.5 |
| Producer body mass ($M_p$) | 1 |
| Basal natural death rate ($d_0$) | 0.4 |
| Allometric slope ($b$) | -0.25 |
| Basal metabolic rate ($a_x$) | producer = 0, ectotherm = 0.88 |
| Maximum consumption rate ($y$) | producer = 0, ectotherm = 4 |
| Assimilation efficiency ($e_{ij}$) | carnivory = 0.85, herbivory = 0.45 |
| Initial richness ($S$) | [10, 20, 40, 60] |
| Predator-Prey Mass Ratio ($Z$) | [1, 5, 10, 25, 50, 100] |
| Predator interference ($c$) | [0, 1] |
| Initial connectance ($C$) | (0.02, 0.38), N = 32 |
| Environmental stochasticity ($\sigma_e$) | [0.1, 0.2, 0.3, 0.4, 0.5, 0.6] |
| Response diversity ($1 - \rho$) | [0, 0.25, 0.5, 0.75, 1] |

Table S2: Standardised estimates from the Structural Equation Model displayed in main text (Fig. 2).

| Response | Predictor | Std estimate |
|---|---|---|
| Population stability | Richness | -0.04 |
| | Avg interaction strength | 0.02 |
| | Avg trophic level | 0.17 |
| | Connectance | -0.02 |
| | Env stochasticity | -0.91 |
| | Predator interference | 0.07 |
| | PPMR | 0.04 |
| | Response diversity | -0.07 |
| Portfolio effect (Doak) | Richness | 0.18 |
| | Avg interaction strength | -0.05 |
| | Avg trophic level | -0.04 |
| | Connectance | -0.01 |
| | Env stochasticity | 0.05 |
| | Predator interference | 0.04 |
| | PPMR | -0.02 |
| | Response diversity | 0.85 |
| Comp effect (Interaction) | Richness | 0.14 |
| | Avg interaction strength | -0.03 |
| | Avg trophic level | -0.17 |
| | Connectance | -0.16 |
| | Env stochasticity | -0.07 |
| | Predator interference | -0.35 |
| | PPMR | 0.08 |
| | Response diversity | -0.12 |
| Asynchrony | Portfolio effect (Doak) | 0.92 |
| | Comp effect (Interaction) | 0.36 |
| Community stability | Population stability | 0.90 |
| | Asynchrony | 0.57 |

Table S3: Summary statistics of the food-web and stability metrics.

| Type | Metric | N | Median (5%, 95%) |
|---|---|---|---|
| **Stability** | Asynchrony | 46880 | 2.1 (1.4,5.8) |
| | Comp effect (Environment) | 46880 | 0.64 (0.35,1.2) |
| | Comp effect (Interaction) | 46880 | 1.2 (0.98,1.6) |
| | Population stability | 46880 | 5.7 (2.4,21.0) |
| | Portfolio effect (Doak) | 46880 | 1.7 (1.1,4.3) |
| | Community stability | 46880 | 14.0 (5.0,56.0) |
| **Food-web** | Avg interaction strength | 46880 | 0.02 (0.0031,0.071) |
| | Avg omnivory | 46880 | 0.04 (0.0,0.25) |
| | Connectance | 46880 | 0.12 (0.058,0.3) |
| | Max trophic level | 46880 | 2.5 (2.0,3.5) |
| | Richness | 46880 | 13.0 (5.0,26.0) |
| | Avg trophic level | 46880 | 1.4 (1.1,2.1) |

Table S4: Variance Inflation Factors for the Structural Equation Model displayed in main text (Fig. 2). All VIF were inferior to 3, indicating low multicollinearity.

| Response | Term | VIF |
|---|---|---|
| **Population stability** | Richness | 2.7 |
| | Avg interaction strength | 2.0 |
| | Avg trophic level | 1.7 |
| | Connectance | 1.6 |
| | Env stochasticity | 1.0 |
| | Predator interference | 1.0 |
| | PPMR | 1.6 |
| | Response diversity | 1.0 |
| **Portfolio effect (Doak)** | Richness | 2.7 |
| | Avg interaction strength | 2.0 |
| | Avg trophic level | 1.7 |
| | Connectance | 1.6 |
| | Response diversity | 1.0 |
| | Predator interference | 1.0 |
| | PPMR | 1.6 |
| | Env stochasticity | 1.0 |
| **Comp effect (Interaction)** | Richness | 2.7 |
| | Avg interaction strength | 2.0 |
| | Avg trophic level | 1.7 |
| | Connectance | 1.6 |
| | Response diversity | 1.0 |
| | Predator interference | 1.0 |
| | PPMR | 1.6 |
| | Env stochasticity | 1.0 |
| **Asynchrony** | Portfolio effect (Doak) | 1.0 |
| | Comp effect (Interaction) | 1.0 |
| **Community stability** | Population stability | 1.0 |
| | Asynchrony | 1.0 |

Table S5: Variance Inflation Factors for the linear model predicting the relationship between community stability and species richness (Fig. 3). All VIF were inferior to 3, indicating low multicollinearity.

| Term | VIF |
|---|---|
| Richness | 2.7 |
| Response diversity | 1.0 |
| Env stochasticity | 1.0 |
| Connectance | 1.6 |
| Avg trophic level | 1.7 |
| PPMR | 1.6 |
| Avg interaction strength | 2.0 |
| Predator interference | 1.0 |
